# JNK-IN-8, a c-Jun N-terminal kinase inhibitor, improves functional recovery through suppressing neuroinflammation in ischemic stroke

**DOI:** 10.1101/449496

**Authors:** Hongli Tang, Qinxue Dai, Wandong Hong, Kunyuan Han, Danyun Jia, Yunchang Mo, Ya Lv, Hongxing Fu, Jianjian Zheng, Wujun Geng

## Abstract

c-Jun N-terminal kinase (JNK), a mitogen activated protein kinase, is activated in ischemia brain injury and plays an important role in cerebral ischemic injury. Emerging studies demonstrated that JNK-IN-8 (a specific JNK inhibitor) regulates traumatic brain injury through controlling neuronal apoptosis and inflammation. However, the role of JNK-IN-8 in ischemic stroke and the underlying mechanisms of JNK-IN-8 involving neuroprotection remain poorly understood. In the present study, male rats were subjected to tMCAO (transient middle cerebral artery occlusion) followed by treatment with JNK-IN-8, and then the modified improved neurological function score (mNSS), the Foot-fault test and the level of inflammatory cytokines (IL-1β, IL-6 and TNF-α) were assessed. We found that JNK-IN-8-treated rats with MCAO exerted a significant improvement in spatial learning as measured by the improved mNSS, and showed sensorimotor functional recovery as measured by the Foot-fault test. JNK-IN-8 also exerted anti-inflammatory effects as indicated by decreased activation of microglia and the decreased expresson of IL-6, IL-1β and TNF-α. Furthermore, JNK-IN-8 suppressed the activation of JNK and subsequent activation of NF-κB signaling as indicated by the decreased level of phosphorylated JNK (p-JNK) and p65. These data suggest that JNK-IN-8 suppressed neuroinflammation and improved neurological function by inhibiting JNK/NF-κB pathway after ischemic brain injury, thus offering a new target for prevention of ischemic brain injury.

## 1. Introduction

Ischemic stroke is well-known as a leading cause of long-term disability worldwide and countless efforts have been made towards the therapeutic treatment on the disease. Despite current diagnosis and prognosis for ischemic stroke can be facilitated by genetic or transcriptomic biomarkers, it becomes seriously limited in terms of early stroke management [1]. It has been demonstrated widely that neuroinflammation plays an important role in the pathophysiology of ischemic injury stroke in which local neuroinflammation can cause neuronal damage and degeneration [2]. Previous clinical studies reported that the prognosis of stroke can be significantly influenced by systemic inflammation [3] while inhibition on inflammatory responses could decrease brain injury [4]. Therefore, a comprehensive understanding of regulation on inflammatory processes in response to the brain injury is a prerequisite in terms of developing effective treatment for ischemic stroke.

c-Jun N-terminal kinase (JNK), a mitogen activated protein kinase, has been shown to be related to inflammatory processes in many diseases [5, 6]. JNK is considered as an important stress-responsive kinase and JNK signaling is reportedly associated with neuroinflammation, blood–brain barrier (BBB) disruption, and oligodendroglial apoptosis in brain injury [7]. Studies on novel therapeutic targets neuroinflammation and neuropathic pain have considered JNK as a promising candidate due to its regulatory roles in inducing neuroinflammation in vivo and in vitro [8]. A specific JNK inhibitor, JNK-IN-8, is a potent and selective covalent inhibitor of JNK and has been used to investigate the precise role of JNK in many pathways and diseases [9]. In a previous study, a dual NO-donating oxime and c-Jun N-terminal kinase inhibitor is reported to protect cells against cerebral ischemia-reperfusion injury in mice, indicating the use of JNK inhibitor is accessible for investigation on functions of JNK in ischemia injury [10]. Specifically, JNK-IN-8 is reported to significantly suppress tumor growth in vitro and in vivo, directly proving JNK regulating breast cancer tumorigenesis [11]. These studies exhibit the valuable aspect of JNK inhibitors in protecting against molecular occurrence of different diseases, however, the precise effect of JNK-IN-8 on neuroinflammation related to ischemic stroke remains unclear.

In the present study, we aimed to investigate whether the use of JNK-IN-8 can improve functional recovery through suppressing neuroinflammation in ischemic stroke. We used established tMCAO (transient middle cerebral artery occlusion) rat model by treatment with JNK-IN-8. We assessed the modified improved neurological function score (mNSS), the Foot-fault test and the level of inflammatory cytokines (IL-1β, IL-6 and TNF-α). Current data demonstrated that JNK-IN-8 exerted a neuroprotective and anti-inflammatory effects and suppressed the activation of JNK and subsequent activation of NF-κB signaling in ischemic brain injury, proposing a potent treatment for ischemic brain injury.

## 2. Materials and methods

### 2.1 MCAO model

All animal experiments were approved by the Animal Ethics Committee at Wenzhou Medical University. Adult male Sprague-Dawley rats (6-8 weeks of age, 200-250g) were obtained from the Shanghai Laboratory Animal Center (Chinese Academy of Sciences, Shanghai, China) and kept in a specific pathogen-free environment for the duration of the current study. Rats were randomly divided into the sham-operated group (n=5) and experimental group (n=15), which was then subdivided into two subgroups (vehicle and JNK-IN-8 group).

The animals were subjected to MCAO as previously described [12, 13]. Briefly, SD rats were anesthetized by choral hydrate (350 mg/kg) intraperitoneally. The right common carotid artery (CCA), internal carotid artery (ICA) and external carotid artery (ECA) were exposed by a midline cervical incision. The ECA was coagulated and inserted into the ICA through the ECA to occlude the MCA. And two hours later, the suture was withdrawn to allow MCA perfusion. The regional cerebral blood flow was observed to verify the occurrence of ischemia by MCAO, using laser Doppler flowmetry (PeriFlux System 5000, Sweden). The sham control rats underwent the same procedures except the occlusion of the MCA. The temperature was maintained at 37.0°C using a heating pad (Malvern, UK) and the rats were kept on it until the closure of the skin incision.

### 2.2 Cell culture

The murine BV2 microglial cells were obtained from the Cell Bank of Chinese Academy of Sciences (Shanghai, China), and cultured in Dulbecco modified Eagle medium (DMEM) supplemented with 10% fetal bovine serum (FBS) and 1% penicillin as well as streptomycin (Gibco BRL, Grand Island, USA), and were maintained in a humidified incubator at 37 □ with 5% atmosphere. The oxygen-glucose deprivation (OGD) was initiated by exposure of BV2 cells to DMEM without serum or glucose in a humidified atmosphere of 95% nitrogen and 5% CO_2_ for 6 hours.

### 2.3 Drug administration

Rats were given an intraperitoneal injection of vehicle (saline 150 δl and 20% dimethylsulfoxide in PBS) or JNK-IN-8 (Selleck Chemicals, TX, USA; 20 mg/kg dissolved in 20% dimethylsulfoxide) after MCAO. Microglia was treated with 10 mM JNK-IN-8 and then pro-inflammatory cytokines (TNF-α, IL-1β and IL-6) was assessed using qPCR and ELISA.

### 2.4 Behavioral Tests

The modified Neurological Severity Score (mNSS) test was used to measure neurological function as previously described [14]. The test was performed on all rats preinjury and at 1, 3, 7, and 14 days after MCAO. The mNSS is a composite of the motor (muscle status and abnormal movement), sensory (visual, tactile and proprioceptive), and reflex tests. Neurological function was graded on a scale of 0-18, where a score of 0 indicates normal performance and a total score of 18 points indicates maximal deficit. In the severity scores of injury, 1 point is awarded for each abnormal behavior or for lack of a tested reflex; thus, the higher the score the more severe the injury.

The foot-fault test was used to evaluate sensorimotor function before MCAO and at 1, 3, 7, and 14 days after MCAO as previously described [15]. The rats were allowed to walk on a grid. With each weight-bearing step, a paw might fall or slip between the wires. If this occurred, it was recorded as a foot fault. A total of 50 steps were recorded for the right forelimb.

### 2.5 Quantitative real-time PCR (qPCR)

Total RNA was extracted from microglia using Trizol reagent (Invitrogen, CA, USA) following the manufacturer’s instructions. Reverse transcription (RT) was carried out using Prime Script TM Master Mix and oligo-dT primers (Takara, Japan). qPCR was performed using 2×SYBR Green Mix (Vazyme Biotech, China) on Applied Biosystems 7300 real-time PCR system (Applied Biosystems, CA, USA). β-actin were used as references for mRNA. The primer information was shown in Supporting Table. S1.

### 2.6 Western blotting

Total protein was extracted from cerebral cortex or microglia with homogenization in lysis buffer and centrifuged at 12,000 rpm for 15 min. Bicinchoninic acid (BCA) assay (Cell Signaling Technology, Boston, MA) was used to determine the protein concentrations. The western blotting was conducted as previously described [16]. After western blotting, the proteins were incubated with specific primary antibodies against Iba-1 (1:1000, ab178846, Abcam, MA, USA), JNK (1:1000, #9252, CST, MA, USA), p-JNK (1:1000, #9251, CST), P65 (1:1000, #3039, CST), I-kBa (1:1000, #9242, CST) overnight at 4 □, and subsequently detected using the secondary antibody (1:2000; Cell Signaling Technology, Boston, MA). β-actin was used as an internal control.

### 2.7 ELISA assay for TNF-α, IL-1β, and IL-6

The ELISA assay was performed using ELISA kit (ADL, San Diego, USA) according to instructions. Briefly, to quantify TNF-α, IL-1β and IL-6 protein level in brain tissues, 96 well plates coated with indicated antibodies were treated with the addition of the supernatants of brain tissue homogenates (1:20 dilution). After the reaction between enzyme and substrate, the absorbances of samples were measured at 450nm with a microplate reader. Standard curves were applied using diluted standard solutions to provide the comparison for the calculation of rat TNF-α, IL-1β and IL-6 in the samples. All the procedures were repeated for at least three times.

### 2.8 Immunofluorescence staining

The immunofluorescence staining in brain tissues was performed as previously described [17]. Specific primary antibody against Iba-1 (1:1500, ab178846) was used to mark the section at 4°C overnight. Secondary antibody for immunofluorescence was anti-rat Alexa Fluor 488 (1:1000, Invitrogen) and then counterstained with DAPI (ATOM, USA). The samples were observed and analyzed by the LEICA TCS SPE microscope (Leica, Germany) and LEICA software LAS AF, respectively. And the positive cells were statistically counted and plotted.

### 2.9 Statistics

All data are presented as mean ± standard deviation (SD) from at least three separate experiments. The differences between groups were analyzed using Student’s *t* test. Differences were deemed statistically significant at *p* < 0.05.

## 3. Results

### 3.1 JNK-IN-8 enhances functional recovery after stroke

c-Jun N-terminal kinases (JNK) are important stress responsive kinases, and JNK signaling is the shared pathway linking microglia activation, neuroinflammation and cerebral ischemic injury [18, 19]. Previous studies demonstrated that JNK inhibition by specific inhibitor (e.g., IQ-1S and SP600125) contributes to reduce cerebral ischemia-reperfusion injury in mice [20, 21]. JNK-IN-8, an adenosine triphosphate-competitive irreversible pan-JNK inhibitor, exerts a protective role in inhibiting traumatic brain injury and cancer cell grow [22, 23]. Here we investigated whether JNK-IN-8 treatment reduces cerebral ischemic injury and regulates ischemia-induced neuroinflammation using experimental model of ischemic stroke.

To test the effect of JNK-IN-8 on regulating cerebral ischemic injury, the sensorimotor performance of the stroke severity was evaluated by assaying foot-fault and modified neurological severity score (mNSS) after JNK-IN-8 treatment. Rats subjected to MCAO, and then received vehicle or JNK-IN-8 intraperitoneally at 2 h after MCAO. The mNSS and Foot-fault test were carried out prior to the treatment after MCAO, at day 1, 3, 7 and 14 after MCAO.

The mNSS was close to 12 in rats with MCAO (both the vehicle and JNK-IN-8 groups) on day 1 post-MCAO, suggesting that neurological functional deficits were comparable in both groups before treatment (Figure 1A). Significant decrease of mNSS was found over time in the vehicle-treated rats on day 3, 7 and 14 after MCAO, indicating that a spontaneous sensorimotor functional recovery occurred after MCAO. Importantly, JNK-IN-8 treatment resulted in a significant functional recovery on day 3, 7 and 14 after MCAO compared with the vehicle (Figure 1A). JNK-IN-8 treatment also reduced the frequency of forelimb foot-fault occurrence compared with vehicle (Figure 1B). These results demonstrated that JNK-IN-8 improves functional recovery after stroke.

**Figure.1.**
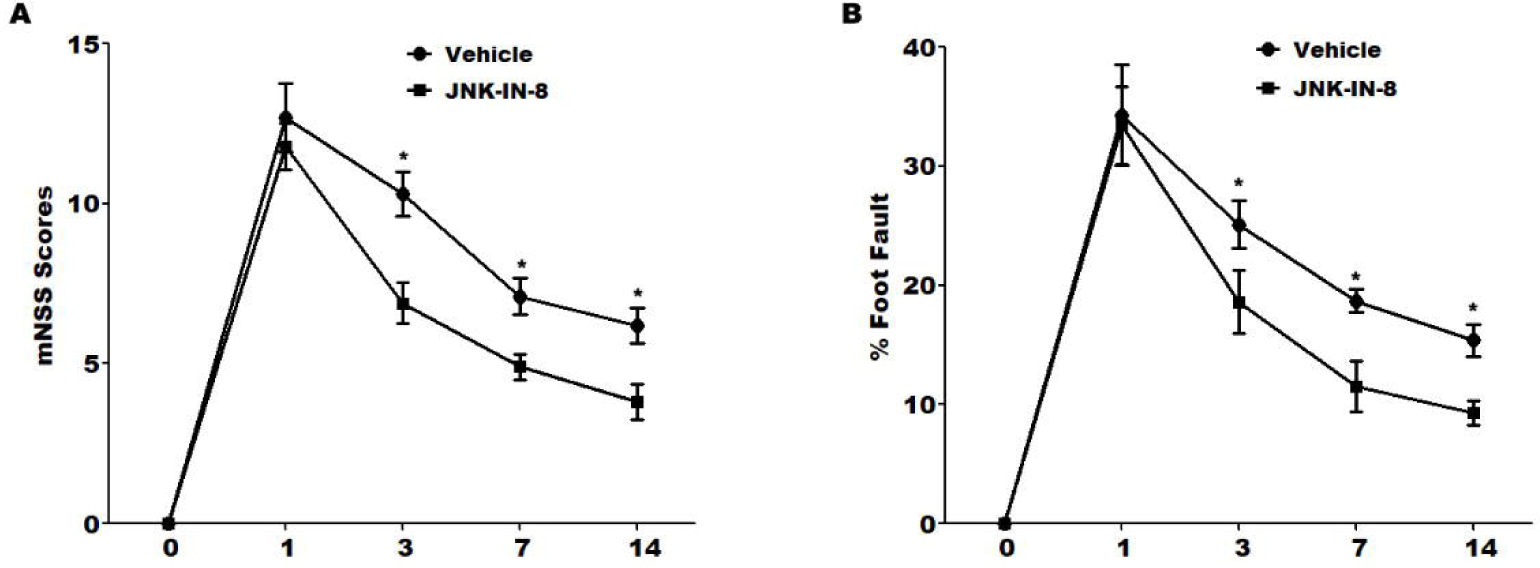
JNK-IN-8 improves functional recovery after stroke. (A) Modified neurological severity scores (mNSS) of rat behavior after MCAO and intraperitoneal injection of vehicle or JNK-IN-8 (20 mg/kg). n = 5. \**P*<0.05. (B) A foot-fault test after MCAO and intraperitoneal injection of vehicle or JNK-IN-8 (20 mg/kg). n = 5. \**P*<0.05. *Single asterisk indicated the significant difference when *P*<0.05.

### 3.2 JNK-IN-8 inhibits microglia activation *in vivo* after stroke

Microglia rapidly responds to cerebral injury through proliferating, changing morphology, and cytokines production [24, 25]. Post-ischemic inflammation is a primary step in the progression of brain ischemia-reperfusion injury [26, 27]. To unveil the potential mechanism of JNK-IN-8 on treating MCAO rats, microglia activation was analyzed by immunofluorescence analysis using Iba-1 antibody. Figure 2A showed that MCAO resulted in a significant microglia activation in ipsilateral cortex at 4 h after MCAO as indicated by the intensive ramified Iba-1-positive staining, which was obviously inhibited by JNK-IN-8 treatment (Figure 2A). To verify the effect of JNK-IN-8 on microglia activation, the protein level of Iba-1 in ipsilateral cortex was assayed by western blot analysis. As shown in Figure 2B, the protein level of the microglial marker Iba-1 was significantly increased in the ipsilateral cortex of MCAO rats compared with the control group of rats, whereas this increase was inhibited by JNK-IN-8 treatment.

**Figure.2.**
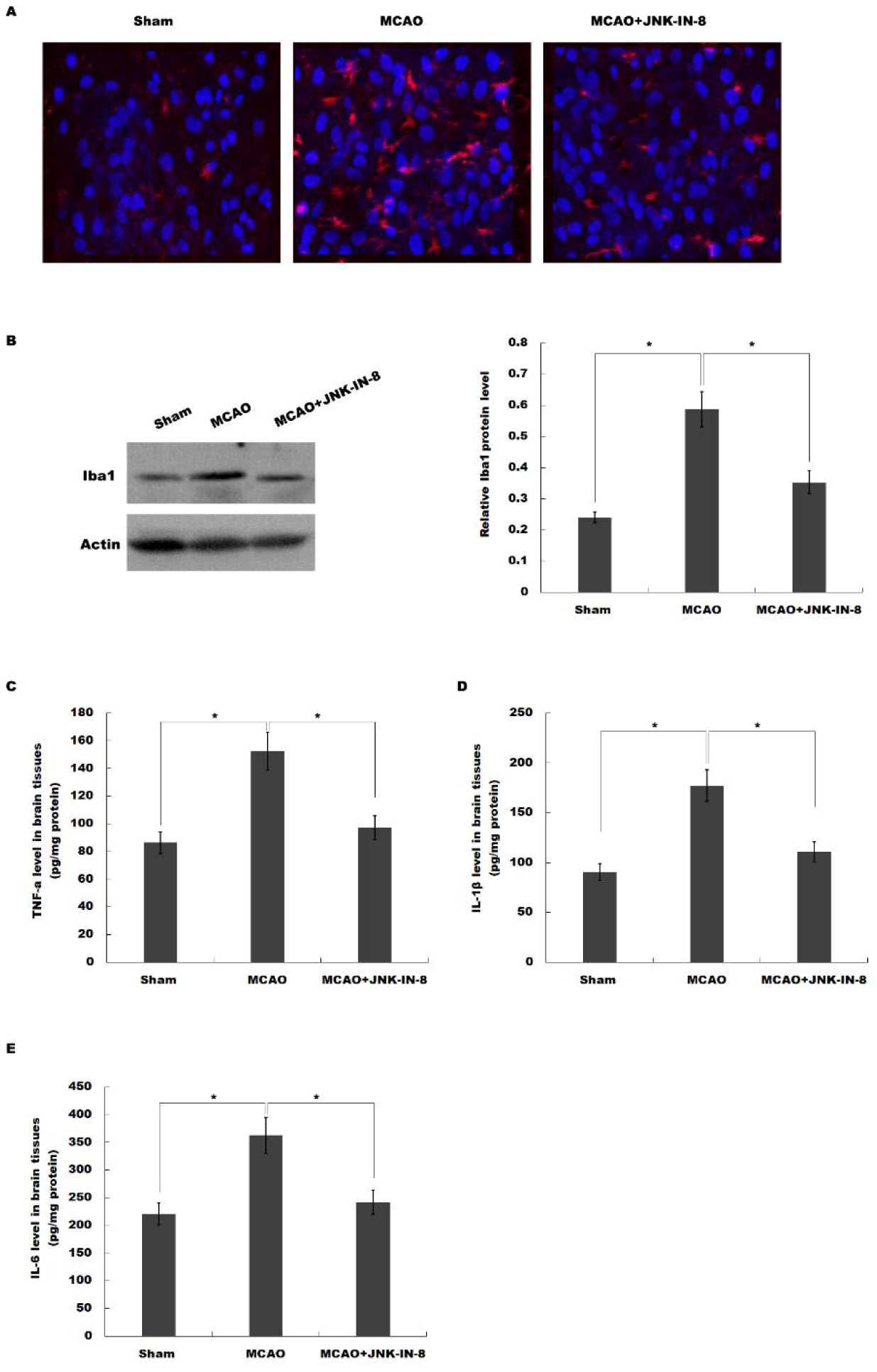
JNK-IN-8 inhibits microglia activation *in vivo* after stroke. (A) Immunofluorescence of microglia cells in ipsilateral cortex after MCAO and intraperitoneal injection of vehicle or JNK-IN-8. (B) Western blot analysis of Iba-1 in ipsilateral cortex after MCAO and intraperitoneal injection of vehicle or JNK-IN-8. (C-E) ELISA analysis of TNF-α (C), IL-1β (D) and IL-6 (E) level in brain tissues after MCAO and intraperitoneal injection of vehicle or JNK-IN-8. \**P*<0.05.

We then assessed the effects of JNK-IN-8 on the production of pro-inflammatory cytokines in brain tissues. ELISA analysis was carried out to assay the protein level of TNF-α, IL-1β and IL-6 in brain tissues. As shown in Figure 2C-E, MCAO resulted in a significant increase of TNF-α, IL-1β and IL-6, whereas JNK-IN-8 treatment inhibited these pro-inflammatory cytokines production. These data suggest that JNK-IN-8 plays important roles in inhibiting cerebral ischemia-induced microglia activation and subsequent neuroinflammation.

### 3.3 JNK-IN-8 inhibits the activation of JNK-NK-κB pathway

We then investigated whether JNK-IN-8 treatment inhibited JNK activation *in vivo* after MCAO. Western blot analysis revealed that brain phosphorylated JNK (p-JNK) levels were increased at 4 h after MCAO, and JNK-IN-8 treatment significantly attenuated brain p-JNK level compared with the vehicle, indicating JNK pathway was activated after MCAO and ischemia-induced JNK activation was inhibited after JNK-IN-8 treatment (Figure 3A and B).

**Figure.3.**
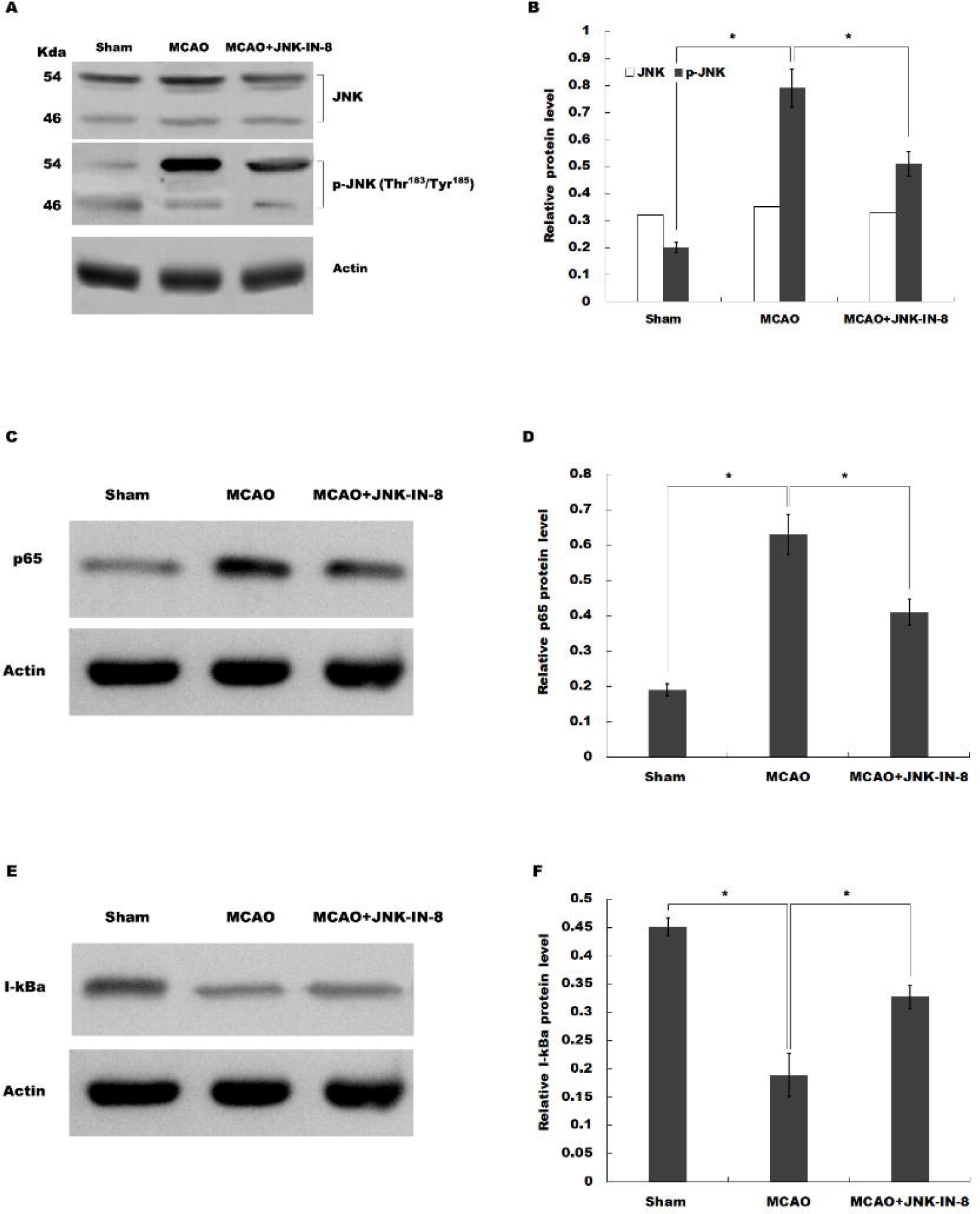
JNK-IN-8 inhibits the activation of JNK-NK-κB pathway. (A and B) Western blot analysis of brain JNK and phosphorylated JNK (p-JNK) levels after MCAO and intraperitoneal injection of vehicle or JNK-IN-8. (C and D) Western blot analysis of brain p65 levels after MCAO and intraperitoneal injection of vehicle or JNK-IN-8. (E and F) Western blot analysis of brain I-kBa levels after MCAO and intraperitoneal injection of vehicle or JNK-IN-8. \**P*<0.05.

NF-κB has been shown to be an important upstream modulator for pro-inflammatory cytokines in microglia [28, 29]. We then examine whether the neuroinflammation-relieving effect of JNK-IN-8 in MCAO rats is related to NF-κB activation. As shown in figure 3C-F, the expression of p65 was increased but ikappaB-alpha (I-KBa) was significantly decreased by ischemic injury, indicating that NF-κB signaling was activated after MCAO. As expected, these changes in NF-κB pathway were reversed by JNK-IN-8 treatment.

### 3.4 JNK-IN-8 inhibits microglia activation and the production of pro-inflammatory cytokines *in vitro*

To verify the role of JNK-IN-8 on microglia activation *in vitro,* the activation of BV2 microglia cells on normoxia or oxygen and glucose deprivation (OGD) with or without JNK-IN-8 treatment was assessed by Cell Counting Kit-8 analysis. As shown in Figure 4A, cell activity of BV2 microglia cells was significantly decreased with JNK-IN-8 treatment under normoxia or OGD, indicating the inhibitory effect of JNK-IN-8 on BV2 microglia activation.

**Figure.4.**
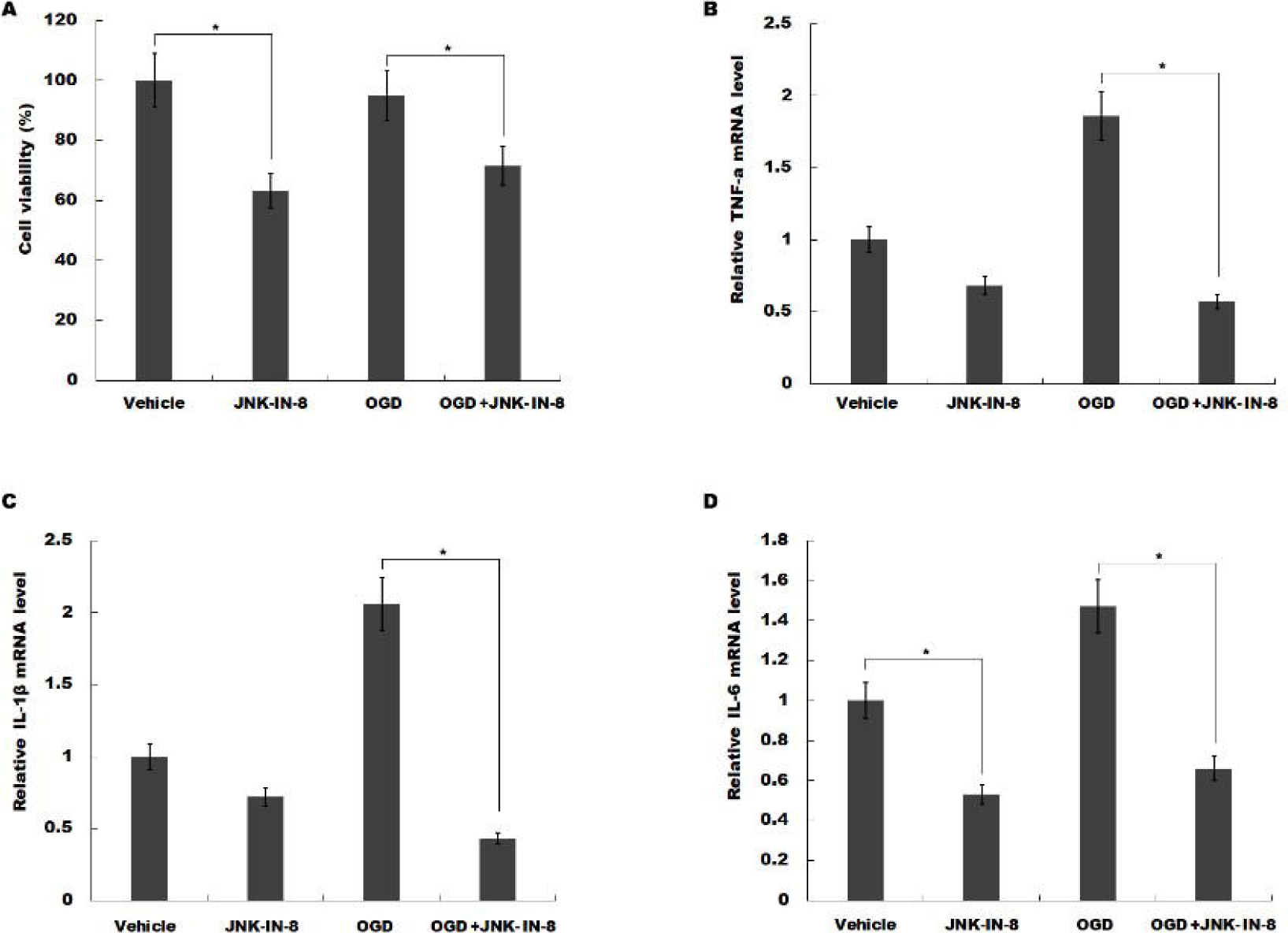
JNK-IN-8 inhibits microglia activation and the production of pro-inflammatory cytokines *in vitro*. (A) JNK-IN-8 treatment (10mM) inhibited BV2 microglia viability determined by Cell Counting Kit-8 kit. (B-D) JNK-IN-8 treatment (10mM) reduced the mRNA levels of TNF-α (B), IL-1|3 (C), and IL-6 (C) in BV2 cells determined by quantitative real-time PCR, BV2 cells under normoxia with OGD groups, BV2 cells subjected to oxygen and glucose deprivation. \**P*<0.05.

We then assessed the effects of JNK-IN-8 on the production of pro-inflammatory cytokines *in vitro.* OGD cellular models were used and then BV2 cells were treated with JNK-IN-8. Figure 4B-D showed that JNK-IN-8 treatment suppressed the mRNA level of TNF-α, IL-1β and IL-6 under normoxia. Furthermore, the expression level of these pro-inflammatory cytokines in BV2 microglia cells with or without JNK-IN-8 treatment on OGD was assayed using qPCR. As shown in Figure 4B-D, TNF-α, IL-1β and IL-6 mRNA levels were increased under OGD, but JNK-IN-8 treatment attenuated OGD-induced upregulation of TNF-α, IL-1β and IL-6 mRNA level. There results suggest that JNK-IN-8 treatment contributes to functional recovery after stroke in rats by suppressing JNK-NF-ΫB pathway, and attenuating microglia activation and pro-inflammatory response.

## 4. Discussion

In this study, we investigated a possible role of JNK-IN-8 in protecting against cerebral ischemia injury in a rat model of MCAO, and determine the precise role of JNK-IN-8 in regulations on stroke. The current data demonstrate that (i) JNK-IN-8 inhibits the activation of JNK-NK-κB pathway, (ii) JNK-IN-8 inhibits microglia activation *in vivo* after ischemic stroke, (iii) JNK-IN-8 inhibits microglia activation and the production of pro-inflammatory cytokines *in vitro*, (iv) JNK-IN-8 improves functional recovery after stroke. These results reveal a potential role of JNK-IN-8 in regulating neuroinflammation, and may provide a novel opportunity to treat ischemic stroke.

Post-ischemic inflammation contributes to the development of neuronal injury and cerebral infarction [30]. However the advances in suitable therapy for the purpose of decreasing neuroinflammation remain limited. The JNK is a mitogen activated protein kinase (MAPK) family member that modulates multiple cellular functions, such as proliferation, apoptosis, and differentiation [31]. JNK signaling is reportedly associated with microglia activation and neuroinflammation [7]. Additionally, JNK mediates the activation of NF-κB signaling, which is an important upstream modulator for pro-inflammatory cytokines in microglia [28, 29]. Therefore, studies on novel therapeutic targets neuroinflammation and neuropathic pain have considered JNK as a promising candidate.

Several synthetic inhibitors of JNK have been described in cerebral ischemia/reperfusion injury, including small molecules SP600125, AS601245 and IQ-1S. Guan *et al* reported that SP600125 treatment inhibits the activation of JNK and provides neuroprotection in ischemia/reperfusion by suppressing neuronal apoptosis [32]. Another study demonstrated that SP600125 attenuated subarachnoid hemorrhage-cerebral vasospasm through a suppressed inflammatory response [33]. IQ-1S releases NO during its oxidoreductive bioconversion and improves stroke outcome in a mouse model of cerebral reperfusion [20]. Noteworthily, different JNK inhibitor exerts diverse physiological properties because of targeting different members of the JNK family. Although currently available candidate JNK inhibitors with high therapeutic potential are identified, the further search for selective JNK inhibitors remains an important task.

In the study, the role of JNK-IN-8 in regulating neuroinflammation and neurological function after stroke was investigated. JNK-IN-8 treatment inhibits stroke-induced microglia activation *in vivo* and *in vitro.* TNF-α, IL-1β and IL-6 levels are increased under OGD, but JNK-IN-8 treatment attenuates OGD-induced upregulation of TNF-α, IL-1β and IL-6 level. There results suggest that JNK-IN-8 treatment contributes to attenuate microglia activation and pro-inflammatory response. As expected, JNK-IN-8 significantly inhibits brain p-JNK level compared with the vehicle, indicating ischemia-induced JNK activation was inhibited by JNK-IN-8. Furthermore, we demonstrated that JNK-IN-8 reduces JNK-mediated activation of NF-κB signaling. More important, MCAO rats treated with JNK-IN-8 exerts a significant improvement in spatial learning and sensorimotor functional recovery as measured by the mNSS and Foot-fault test. Taken together, this study suggests an interesting prospect to target JNK-IN-8 as a potent therapy for ischemic stroke.

## Data accessibility

This article has no additional data.

## Authors’ contributions

H.T., O.D., D.J. and S.L. participated in study design and carried out the molecular laboratory work, acquisition of data, data analysis and interpretation, and drafting of the manuscript. J.T. and Y.L. helped to draft the manuscript. W.G. contributed to the conception and design of the study and final approval of the submitted version. All authors read and approved the final manuscript. The authors declare that there are no conflicts of interest regarding the publication of this article.

## Acknowledgments

The project was supported by the National Natural Science Foundation of China (No.81774109,No.81603685,No. 81704180), Zhejiang Provincial Department of Education (No. Y201839270), and Wenzhou Municipal Science and technology Bureau (No. Y20170023,No. Y20170164,No. Y20170141).

